## Supplementary material for "Controlling microbial co-culture based on substrate pulsing can lead to stability through differential fitness advantages": S1_file

### Supplementary File 1: Extended Model description

#### Monod-type ODE model construction

As described in the main manuscript, a simplified cybernetic mathematical framework for a single strain comprising different metabolite consumption and production states was constructed first for each organism. Each single-strain cybernetic model was used as a kernel section of a co-culture population computational framework. A Monod base model was used for the individual strain rate equations. For this construction, let us first establish a common biochemical reaction for any substrate $S$ that can be consumed by the biomass $X$ at a given metabolic state as:

$$XX + Y_{\frac{s}{\sigma}} S \xrightarrow{r_\sigma} X + Y_{\frac{x}{\sigma}} X + Y_{\frac{a}{\sigma}} A + ... + Y_{\frac{n}{\sigma}} N \tag{1}$$

$$\tag{2}$$

The latter biochemical reaction model allows to construct a stoichiometric vector $\phi$ for $S$ consumption such that the rate of change for any component in the media $M_i$ (including $X$ and $S$,$i = X, S, A, ..., N$) can be defined by the product of reaction rate $r_\sigma$ and $\phi$ vector as follows:

$$\frac{dM}{dt} = \phi r_\sigma = \begin{bmatrix} Y^{\sigma}_{\frac{x}{x}} \\ Y^{\sigma}_{\frac{s}{x}} \\ Y^{\sigma}_{\frac{a}{x}} \\ ... \\ Y^{\sigma}_{\frac{n}{x}} \end{bmatrix} r_\sigma \tag{3}$$

With this in mind, the model can then be extended to a stoichiometric Matrix where we can define each column as a different $\phi$ for each known growth rate derived from the consumption of other substrates, which in turn can be associated with a specific metabolic state(e.g., the consumption of $A, ..., N$). This extension then contains the known behavioral capabilities of the cell within the model $\Phi = [\phi_s, \phi_a, ..., \phi_n]$.

$$\frac{dM}{dt} = \Phi \vec{r} = \begin{bmatrix} Y^{\sigma}_{\frac{x}{x}} & Y^{\alpha}_{\frac{x}{x}} & ... & Y^{\nu}_{\frac{x}{x}} \\ Y^{\sigma}_{\frac{s}{x}} & Y^{\alpha}_{\frac{s}{x}} & ... & Y^{\nu}_{\frac{s}{x}} \\ Y^{\sigma}_{\frac{a}{x}} & Y^{\alpha}_{\frac{a}{x}} & ... & Y^{\nu}_{\frac{a}{x}} \\ ... & ... & ... & ... \\ Y^{\sigma}_{\frac{n}{x}} & Y^{\alpha}_{\frac{n}{x}} & ... & Y^{\nu}_{\frac{n}{x}} \end{bmatrix} \begin{bmatrix} r_\sigma \\ r_\alpha \\ ... \\ r_\nu \end{bmatrix} \tag{4}$$

Each $\phi$ vector also contains the information for its effect on all components in the system. If a particular metabolite is not produced or consumed in a specific metabolic state, its yield ($Y$) is set to 0. For our particular purpose, we will put that for a metabolic state for a single primary substrate $(j...j+n)$ consumption, the consumption rate $q_{j....j+n}$ is given by a simplified Monod-type equation such as:

$$q_j = \frac{q_j^{max} M_j}{K_j + M_j} \tag{5}$$

We can then rewrite Eq. 4 in a general form as follows:

$$\frac{dM_i}{dt} = Y_{\frac{i}{\sigma}}^{\sigma} \frac{q_\sigma^{max} M_\sigma}{K_\sigma + M_\sigma} X + Y_{\frac{i}{\alpha}}^{\alpha} \frac{q_\alpha^{max} M_\alpha}{K_\alpha + M_\alpha} X + ... + Y_{\frac{i}{\nu}}^{\nu} \frac{q_\nu^{max} M_\nu}{K_\nu + M_\nu} X \tag{6}$$

This general form can then be reduced for each metabolite as some yields are 0 for a particular Substrate $j$. In this work, we used this equation to construct the mathematical model to describe the growth, consumption, and production behavior of *E.coli* and *S.cerevisiae* strains. In this simplified model, the biomass $X$ can interact by consuming and producing three different external metabolites, Glucose (GLC, $G$), Acetate (ACE, $A$), and Ethanol (ETH, $E$). These metabolites were chosen as they are the most relevant for the metabolism of both strains when cultured in minimal media with GLC as the only carbon source. The $\Phi$ matrix-related equations constructed were then set to be the following:

$$\frac{dM}{dt} = \Phi \vec{r} = \begin{bmatrix} Y_{\frac{x}{g}}^{\gamma o} & Y_{\frac{x}{g}}^{\gamma f} & Y_{\frac{x}{a}}^{\alpha} & Y_{\frac{x}{e}}^{\epsilon} \\ -1 & -1 & 0 & 0 \\ 0 & Y_{\frac{a}{g}}^{\gamma f} & -1 & 0 \\ 0 & Y_{\frac{e}{g}}^{\gamma f} & 0 & -1 \end{bmatrix} \begin{bmatrix} q_{\gamma o} X \\ q_{\gamma f} X \\ q_\alpha X \\ q_\epsilon X \end{bmatrix} \tag{7}$$

Where $q_{\gamma o}$ $q_{\gamma f}$ are the glucose consumption rates for the oxidative and fermentative pathways, respectively, and $q_\alpha$ and $q_\epsilon$ are the consumption rates for acetate and ethanol. In this work, the rates were also set to be affected by a general metabolism inhibition proportioned by the external accumulation of the substrates as follows:

$$q_j = \frac{q_j^{max} M_j}{K_j + M_j} H^- \tag{8}$$

where:

$$H^- = \Pi_i^n \frac{1}{1 + \frac{M_i}{I_i}} \tag{9}$$

Finally, with the addition of a first-order death rate, the equations used in the present work can be expressed as follows:

$$\frac{dG}{dt} = \left[ -\frac{q_{\gamma o}^{max} G}{K_{\gamma o} + G} - \frac{q_{\gamma f}^{max} G}{K_{\gamma f} + G} \right] X H^- \tag{10}$$

$$\frac{dA}{dt} = \left[ Y_{\frac{a}{g}}^{\gamma f} \frac{q_{max}^{\gamma f} G}{K_{\gamma f} + G} - \frac{q_{max}^{\alpha} A}{K_\alpha + A} \right] X H^- \tag{11}$$

$$\frac{dE}{dt} = \left[ Y_{\frac{e}{g}}^{\gamma f} \frac{q_{max}^{\gamma f} G}{K_{\gamma f} + G} - \frac{q_{max}^{\epsilon} E}{K_\epsilon + E} \right] X H^- \tag{12}$$

$$\frac{dX}{dt} = \left[ Y_{\frac{x}{g}}^{\gamma o} \frac{q_{max}^{\gamma o} G}{K_{\gamma o} + G} + Y_{\frac{x}{g}}^{\gamma f} \frac{q_{max}^{\gamma f} G}{K_{\gamma f} + G} + Y_{\frac{x}{a}}^{\alpha} \frac{q_{max}^{\alpha} A}{K_\alpha + A} + Y_{\frac{x}{e}}^{\epsilon} \frac{q_{max}^{\epsilon} E}{K_\epsilon + E} \right] X H^- - K_d X) \tag{13}$$

*E.coli* and *S.cerevisiae* models were constructed in this work, approximated from their observed behavior in axenic cultures as presented in the main manuscript.

**General description for the Cybernetic framework**

The cybernetic modeling approach was used to extend the characterization of the behavior of the single cultures of *S.cerevisiae* and *E. coli* during the batch cultures and continuous culturing processes as it can render the allocation of cell resources on several metabolic options. The cybernetic variables represent the expression of the metabolic machinery related to a particular substrate metabolism, a so-called "representative enzyme." This representative enzyme encompasses all the essential enzymes, co-factors, and other resources necessary for the metabolic reactions regarding a particular substrate consumption and fate [1]. The cybernetic approach has been previously used to address the diauxic behavior in *Kleibsiela oxytoca* [2], and more recently to derive full dynamic models into the metabolic fluxes across several microorganisms [3–5] and even mammalian cells [3,6]. A full description of the cybernetic approach can be found in Ramkrishna et al. reports [1,2,4–6]. In this work, it was first assumed that each phenotype could be simplified as the consumption of one or more substrates ($M_s$) catalyzed by a critical enzyme ($\Psi_s$) for the production of biomass $X$ and other products ($M_p$) [2]. $\Psi_s$ represents the set of all enzymes governing the kinetics of this specific phenotype. $\Psi_s$ synthesis is induced by the presence of the specific substrate or metabolite $M_s$. This simplified model can be written as:

$$X + M_s \xrightarrow{\Psi_s} (1 + Y_{x/s})X + Y_{p/s}M_p + ... \tag{14}$$

$$X + M_s \xrightarrow{M_s} X' + \Psi_s \tag{15}$$

Where $X'$ represents the biomass excluding the *critical enzyme* $\Psi_s$. These two reactions can be described by known kinetic equations such as the Michaelis-Menten model for enzymatic catalysis. In this work, these kinetic equations are derived from the previously shown mass balance model:

$$\frac{dm_s}{dt} = \frac{q_s \psi_s M_s X}{K_s + M_s} H^- \tag{16}$$

$$\frac{d\psi_s}{dt} = \epsilon_s^c + \frac{\epsilon_s^i M_s X}{K_s' + M_s} H^- - \delta_s \psi_s - \mu \psi_s \tag{17}$$

Where $\epsilon_s^c$ and $\epsilon_s^i$ are the production rate constants for the enzyme for its constitutive and inducible expression, respectively. While $\delta_s$ is the decay constant of the enzyme. These parameters have been previously approximated for various microorganisms and cell lines including *E.coli* and *S.cerevisiae* [1,2,5,7](see Fig.1). $\psi_s$ is the specific concentration of the enzyme $\Psi_s$ such that $\psi_s X$ is the total concentration of this enzyme. $\epsilon_s^c + \epsilon_s^i$ gives the maximum synthesis rate for this enzyme. The cybernetic approach solves the difficulty of calculating $\psi_i$ by assuming that the maximum quantity of enzyme defines the maximum rate. Therefore:

$$q_s^{max} = q_s \psi_s^{max} \tag{18}$$

$$\psi_s^{max} = \frac{\epsilon_s^c + \epsilon_s^i}{u^{max} + \delta_s} \tag{19}$$

the enzyme concentration value can be substituted by a relative enzyme value respective to the maximum enzyme concentration as:

$$q_g \psi_g = q_g^{max} \left[ \frac{\psi_g}{\psi_g^{max}} \right] \tag{20}$$

Finally, the cybernetic modeling introduces the regulation of the inhibition/activation of enzyme expression and repression/induction of enzyme activity by the introduction of the variables $\upsilon$ and $\nu$, which regulate enzyme synthesis $\frac{d\psi_g}{dt}$ and activity $\frac{dm_g}{dt}$ along with the model.

$$\frac{d\Psi_g}{dt} = \upsilon_g \frac{d\psi_g}{dt} \quad (0 < \upsilon_g < 1 \; ; \; \sum_{j=g}^{e} \upsilon_j = 1) \tag{21}$$

$$\frac{dM_g}{dt} = \nu_g \frac{dm_g}{dt} \quad (0 \leq \nu_g \leq 1) \tag{22}$$

The cybernetic variables $\upsilon$ and $\nu$ are calculated by matching law equations constructed for specific metabolic objectives. In the case of this work, the growth rate was selected as the metabolic objective, which in turn represents a fitness index.

Cybernetic variables can then be understood as the comparison between the fitness advantage return for each reaction driven by each $\Psi_{\iota\ldots\omega}$, which can be used to regulate its participation in the cellular metabolism at any time. The equations used for these cybernetic variables are the following:

$$\upsilon_\iota = \frac{\mu_\iota}{\sum_{j=\iota}^{\omega} \mu_j} \tag{23}$$

$$\nu_\iota = \frac{\mu_\iota}{max(\mu_{\iota\ldots\omega})} \tag{24}$$

where $\mu_g$ represents the growth rate supported by each reaction driven by each $\Psi_{\iota\ldots\omega}$ in this case by the consumption of glucose $\Psi_g$, acetate $\Psi_a$ and ethanol $\Psi_e$. This way, the dynamic distribution of its participants can be calculated to describe the metabolic and physiological behavior $\Phi$ given a metabolic reaction network [1,2,4,8]. The approach allows to approximate the current phenotype by describing the metabolite content ($M_{\iota\ldots\omega}$), enzymatic content ($\Psi_{\iota\ldots\omega}$) and its functional relationship given by the regulation ($\upsilon_{\iota\ldots\omega}$ and $\nu_{\iota\ldots\omega}$).

#### Model Parametrization and Approximation values

**Table 1. Cybernetic model parametrization for the enzymatic rate equation for both strains**

| Parameter | Value | Reference |
|---|---|---|
| $\epsilon^c_{coli}$ | 0.01 | [1,5] |
| $\epsilon^i_{coli}$ | 1 | [1,5] |
| $\delta_{coli}$ | 0.05 | [1,5] |
| $\epsilon^c_{sacc}$ | 0.1 | [2,7] |
| $\epsilon^i_{sacc}$ | 0.2 | [2,7] |
| $\delta_{coli}$ | 1 | [2,7] |

**Table 2. Parameters approximated from data for each organism model.**

| Parameter | *E. coli* | *S. cerevisiae* |
|---|---|---|
| $\mu^{ox}_{max}$ | 0.234 | 0.221 |
| $\mu^{ferm}_{max}$ | 0.437 | 0.299 |
| $\mu^{a}_{max}$ | 0.077 | 0.017 |
| $\mu^{e}_{max}$ | — | 0.080 |
| $q^{ox}_{max}$ | -1.745 | -3.175 |
| $q^{ferm}_{max}$ | -2.640 | -1.648 |
| $q^{a}_{max}$ | -0.630 | -0.058 |
| $q^{e}_{max}$ | — | -0.276 |
| $K^{ox}_s$ | 1.000 | 1.486 |
| $K^{ferm}_s$ | 0.112 | 0.082 |
| $K^{a}_s$ | 0.108 | 0.007 |
| $K^{e}_s$ | — | 0.055 |
| $K_d$ | 0.0053 | 0.014 |
| $Y^{ox}_{x/s}$ | 0.134 | 0.070 |
| $Y^{ferm}_{x/s}$ | 0.165 | 0.182 |
| $Y^{ferm}_{a/s}$ | 0.062 | 0.075 |
| $Y^{ferm}_{e/s}$ | 0.000 | 0.387 |
| $Y_{x/a}$ | 0.123 | 0.285 |
| $Y_{x/e}$ | — | 0.287 |

*All inhibition constants were found to be not relevant for present model calculations and were given values of 9999.

**Table 3. Weighted SSE and Willmott's index, and their normalized agreement values as a qualifying measure of the model prediction error for model approximation to various initial GLC concentration experiments.**

| Strain | SSE | WLM | nSSE | nWLM |
|---|---|---|---|---|
| *E. coli* | 1.0527 | 1.6120 | 0.1755 | 0.2687 |
| *S. cerevisiae* | 2.3472 | 0.9689 | 0.3912 | 0.1615 |

### References


1. Ramkrishna D, Song HS. Dynamic Models of Metabolism: Review of the Cybernetic Approach. Bioengineering,Food, and Natural Products. 2012;58(4):986–997. doi:10.1002/aic.
2. Kompala DS, Ramkrishna D, Jansen NB, Tsao GT. Investigation of bacterial growth on mixed substrates: Experimental evaluation of cybernetic models. Biotechnology and Bioengineering. 1986;28(7):1044–1055. doi:https://doi.org/10.1002/bit.260280715.
3. Martinez JA, Bulte DB, Contreras MA, Palomares LA, Ramirez OT. Dynamic Modeling of CHO cell Metabolism Using the Hybrid Cybernetic Approach With a Novel Elementary Mode Analysis Strategy. Frontiers in Bioengineering and Biotechnology. 2020;(8):279.
4. Varner J, Ramkrishna D. Metabolic Engineering from a Cybernetic Perspective Aspartate Familiy of Amino Acids. Metabolic engineering. 1999;1(MT980104):88–116. doi:1096-7176/99.
5. Song HS, Ramkrishna D, Pinchuk GE, Beliaev AS, Konopka AE, Fredrickson JK. Dynamic modeling of aerobic growth of Shewanella oneidensis. Predicting triauxic growth, flux distributions, and energy requirement for growth. Metabolic Engineering. 2013;15:25–33. doi:https://doi.org/10.1016/j.ymben.2012.08.004.
6. Aboulmouna L, Raja R, Khanum S, Gupta S, Maurya MR, Grama A, et al. Cybernetic modeling of biological processes in mammalian systems. Current Opinion in Chemical Engineering. 2020;30:120–127. doi:https://doi.org/10.1016/j.coche.2020.100660.
7. Song HS, Morgan JA, Ramkrishna D. Systematic development of hybrid cybernetic models: Application to recombinant yeast co-consuming glucose and xylose. Biotechnology and Bioengineering. 2009;103(5):984–1002. doi:https://doi.org/10.1002/bit.22332.
8. Ramkrishna D, Song HS. Analysis of Bioprocesses. Dynamic Modeling is a Must. materialstoday:proceedings. 2016;3:3587–3599.
