## Supplementary material for "Controlling microbial co-culture based on substrate pulsing can lead to stability through differential fitness advantages": S2_file

### Supplementary File 2: Simplified MONKS simulation workflow

The MONod-type Co-culture Kinetic Simulation toolbox (MONCKS) is part of the MiPI Bioprocess Modeling and Simulation toolbox (mBMStoolbox) available at .... The former toolbox starts by taking the cybernetic model parameters for two strains, i.e., *E. coli* and *S. cerevisiae*, and using them in a numerical integration calculation with a small $\Delta t$ (i.e., 0.01 h) to calculate their interaction with a common external metabolite pool which simulates the environmental conditions of the bioreactor. Each $\Delta t$, it is possible to add a flux of metabolites to simulate a constant flow or pulses. MONCKS use in the present work was comprised of a series of functions that allow parametrizing the initial conditions of the bioprocess (`BMS_MONCKS_pBioreactor_xlsx`), read and parametrize two strains forming the co-culture system (`BMS_MONCKS_rModels`), perform the numerical simulation of co-culture growth during the bioprocess (`BMS_MONCKS_cSimulation`), read Experimental data sets (`BMS_MONCKS_rExpData_mat`) and compare the fraction of events experimental data to the simulations to assess the simulation quality (`BMS_MONCKS_qSimulation_Xfrac`). The simplified example pipeline in MATLAB can be written as follows:

```
%%%␣MONCKS␣Pipeline␣example␣for␣the␣S.␣cerevisiae␣and␣E.coli␣co-culture␣used␣in␣this␣work
[BMS_BioPar]=BMS_MONCKS_pBioreactor_xlsx('MONCKS_HiFreq.xlsx');
%%%␣File␣in␣brackets␣can␣be␣changed␣for,i.e.,'␣MONCKS_LowFreq.xlsx',␣'MONCKS_Cont.xlsx'
[BMS_AModel,BMS_BModel,BMS_BioPar]=BMS_MONCKS_rModels(BMS_BioPar);
[BMS_BioData,BMS_ASimData,BMS_BSimData]=BMS_MONCKS_cSimulation(BMS_AModel,BMS_BModel,BMS_BioPar);
[BMS_ExpPar,BMS_ExpData]=BMS_MONCKS_rExpData_mat('MONCKSExpDataHifreq');
%%%␣File␣in␣brackets␣can␣be␣changed␣for,i.e.,'␣MONCKS_LowFreq.xlsx',␣'MONCKS_Cont.xlsx'
[BMS_BioPar,BMS_QlyData]=BMS_MONCKS_qSimulation_Xfrac(BMS_BioPar,BMS_BioData,BMS_ExpData);
```

The simulation pipeline starts by reading a .xlsx file with a tab named `Bioreactor` containg the initial fermentation data (Fig. 1). Where `BIOa` and `BIOb` refers to the first and second `.mat` cybernetic model parameter files, respectively. `To`, `Tf` and `dt` are the initial, final and delta times, respectively. `Feed type` can be set in "Discontinuous" and "Continuous" modes, referring to pulses on Feeding times or pulses on Feed concentrations, respectively. `Feed start` is in hours, `Dilution rate` in 1/h, and the `Volume` in L. the `Feed function` used in this work was a "square" wave function, and the `Period Frequency` and `On Time Fraction` are set for simulation in this file as can be seen on the example of Figure 1. The final part of this file contains the vectors of concentrations for the diverse metabolites, including the starting biomass of both microorganisms, for the Feed and the Initial Concentration (Fig. 1).

After reading the initial bioprocess file, the pipeline continues with the reading of the model parameters for the defined microorganisms. These files are `.mat` files containing the structure `BMS_Model`, which includes the main parameters for the cybernetic simulation of the behavior of a particular strain. An example can be seen in the Figure 2:

After acquiring the model parameters for each strain, the pipeline proceeds to numerically simulate the co-culture and the bioreactor environmental conditions according to the simplified algorithm shown in Figure 3.

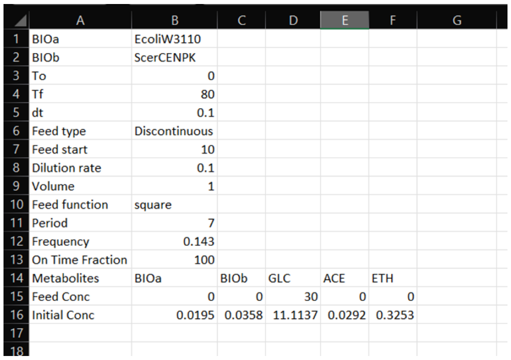

| | A | B | C | D | E | F | G |
|---|---|---|---|---|---|---|---|
| 1 | BIOa | EcoliW3110 | | | | | |
| 2 | BIOb | ScerCENPK | | | | | |
| 3 | To | 0 | | | | | |
| 4 | Tf | 80 | | | | | |
| 5 | dt | 0.1 | | | | | |
| 6 | Feed type | Discontinuous | | | | | |
| 7 | Feed start | 10 | | | | | |
| 8 | Dilution rate | 0.1 | | | | | |
| 9 | Volume | 1 | | | | | |
| 10 | Feed function | square | | | | | |
| 11 | Period | 7 | | | | | |
| 12 | Frequency | 0.143 | | | | | |
| 13 | On Time Fraction | 100 | | | | | |
| 14 | Metabolites | BIOa | BIOb | GLC | ACE | ETH | |
| 15 | Feed Conc | 0 | 0 | 30 | 0 | 0 | |
| 16 | Initial Conc | 0.0195 | 0.0358 | 11.1137 | 0.0292 | 0.3253 | |
| 17 | | | | | | | |
| 18 | | | | | | | |

**Fig 1.** Example of the Bioreactor tab on the .xlsx file for $BMS\_MONCKS\_pBioreactor\_xlsx$ function to read the Bioproccess initial parameters.

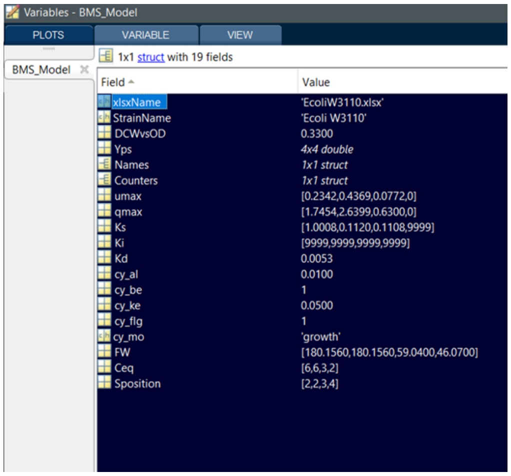


| Field | Value |
|---|---|
| xlsxName | 'EcoliW3110.xlsx' |
| StrainName | 'Ecoli W3110' |
| DCWvsOD | 0.3300 |
| Yps | 4x4 double |
| Names | 1x1 struct |
| Counters | 1x1 struct |
| umax | [0.2342,0.4369,0.0772,0] |
| qmax | [1.7454,2.6399,0.6300,0] |
| Ks | [1.0008,0.1120,0.1108,9999] |
| Ki | [9999,9999,9999,9999] |
| Kd | 0.0053 |
| cy_al | 0.0100 |
| cy_be | 1 |
| cy_ke | 0.0500 |
| cy_flg | 1 |
| cy_mo | 'growth' |
| FW | [180.1560,180.1560,59.0400,46.0700] |
| Ceq | [6,6,3,2] |
| Sposition | [2,2,3,4] |

**Fig 2.** Example of the $BMS\_Model$ structure in the .mat file containing a specific organism and strain model.

Finally, the pipeline reads and compares selected experimental data to the. In this case, we used the fraction of events as the quality reference for the modeling approximation. All the functions and data sets can be found in the repositories at ....

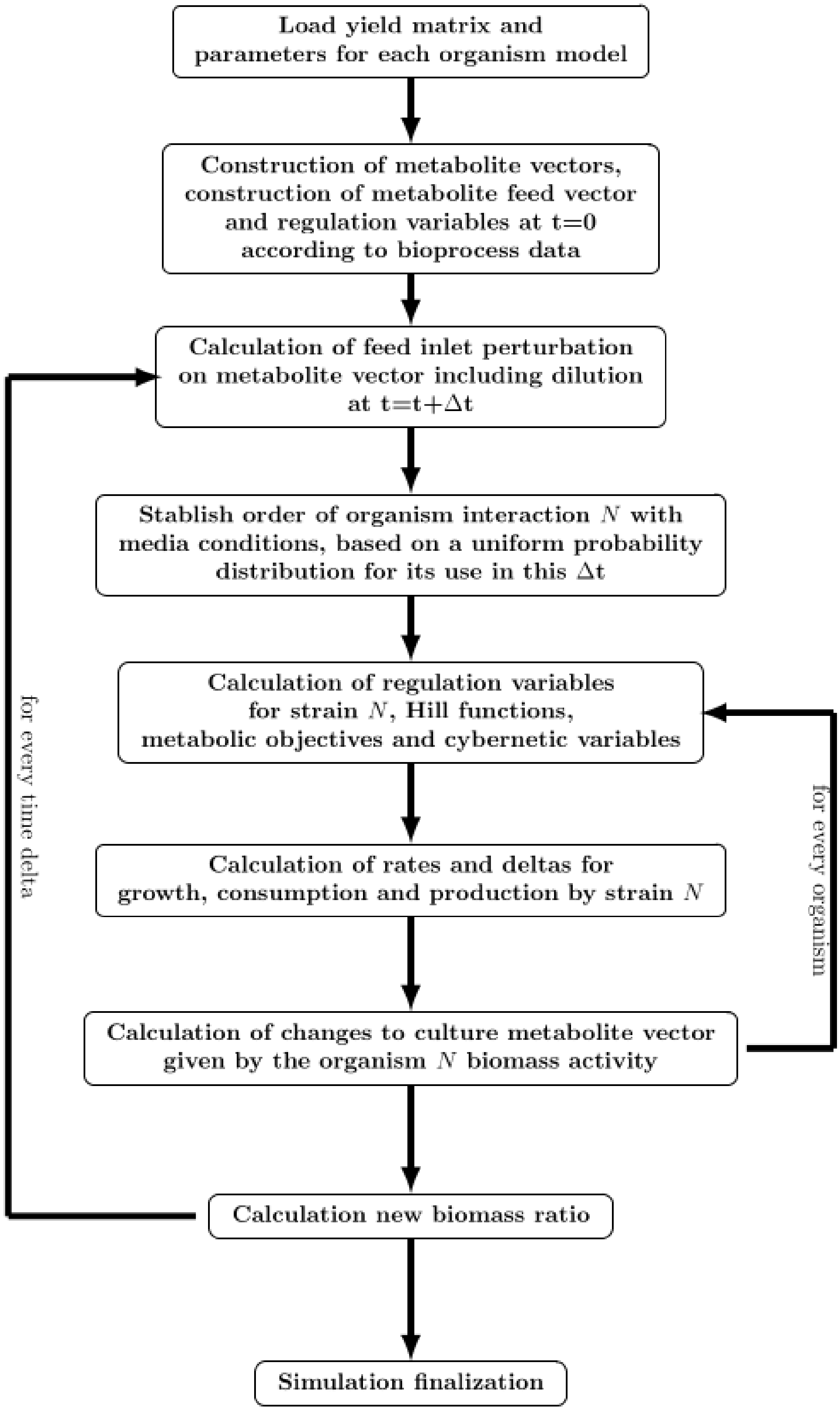


**Fig 3.** Simplified algorithm steps diagram used in MONCKS framework used for simulating the co-culture experiments
