## Supplementary material for "Controlling microbial co-culture based on substrate pulsing can lead to stability through differential fitness advantages": S3_file

### Supplementary File 3: Simplified FC data treatment workflow

Flow cytometry data treatment was performed with the MiPI Flow Cytometry Analysis toolbox (mFCAtoolbox) available at .... This toolbox is a collection of functions for performing quality control and assesment of data, automated gating and clustering of events and statistical analysis on different populations. In this work it was used to address the proportion of *S. cerevisiae* and *E. coli* cells, as well as their statistical properties such as probability density functions for the Front Scattering Channel Area (FSCA), Side Scattering Channel Area (SSCA) and the Flourescence Light sensor 1 Area (FL1A, measuring GFP). An example pipeline used in this work is presented below:

```
␣␣␣%%␣Reading␣and␣organizing␣Data
␣␣␣␣␣␣␣␣FCSFiles=FCA_rFolder(Folder,'online');
␣␣␣␣␣␣␣␣FCSData=FCA_rData(FCSFiles,'usual+h');
␣␣␣␣␣␣␣␣[FCSFiles,FCSData]=FCA_tSelection(FCSFiles,FCSData,to,tf);

␣␣␣␣%%␣Data␣preprocessing
␣␣␣␣␣␣␣␣[FCSFiles,FCSData]=FCA_cData(FCSFiles,FCSData);
␣␣␣␣␣␣␣␣[FCSFiles,FCSData]=FCA_FindDoublets(FCSFiles,FCSData);
␣␣␣␣␣␣␣␣disp('Eliminating␣Doublets␣for␣further␣analysis␣...')
␣␣␣␣␣␣␣␣[FCSFiles,FCSData]=FCA_Selection(FCSFiles,FCSData,'Doublet',0);

␣␣␣␣%%␣Data␣Analysis
␣␣␣␣␣␣␣␣[FCSFiles,FCSData,FCSStats]=FCA_Statistics(FCSFiles,FCSData,[{'FSCA'},{'SSCA'},...
␣␣␣␣␣␣␣␣{'FL1A'}],'type','Logarithmic','distribution','Normal');

␣␣␣␣%%␣Gating␣and␣individual␣stats
␣␣␣␣␣␣␣␣[FCSFiles,FCSData]=FCA_LineGate(FCSFiles,FCSData,'FSCA',ESGate);
␣␣␣␣␣␣␣␣[Ec_FCSFiles,Ec_FCSData]=FCA_Selection(FCSFiles,FCSData,'gates',0);
␣␣␣␣␣␣␣␣[Sc_FCSFiles,Sc_FCSData]=FCA_Selection(FCSFiles,FCSData,'gates',1);
␣␣␣␣␣␣␣␣[Ec_FCSFiles,Ec_FCSData,Ec_FCSStats]=FCA_Statistics(Ec_FCSFiles,Ec_FCSData,[{'FSCA'},{'SSCA'},{'FL1A'}
␣␣␣␣␣␣␣␣[Sc_FCSFiles,Sc_FCSData,Sc_FCSStats]=FCA_Statistics(Sc_FCSFiles,Sc_FCSData,[{'FSCA'},{'SSCA'},{'FL1A'}
␣␣␣␣%%␣Quantification␣of␣aggregates
␣␣␣␣␣␣␣␣[Sc_FCSFiles,Sc_FCSData,Sc_FCSStats]=FCA_GMValueClusters(Sc_FCSFiles,Sc_FCSData,Sc_FCSStats,'FSCA');
␣␣␣␣␣␣␣␣[Ec_FCSFiles,Ec_FCSData,Ec_FCSStats]=FCA_GMValueClusters(Ec_FCSFiles,Ec_FCSData,Ec_FCSStats,'FSCA');
␣␣␣␣␣␣␣␣disp('␣␣␣')
```

All Pipelines start with a reading and organizing Data section, where `FCA_rFolder` lists and reads all the meta data of every `.fcs` file found in the working directory folder. This list is pipelined to `FCA_rData` which performs the parsing and initial organization of the full data for every data point in the set. After this, `FCA_tSelection` helps selecting the time frame we want to analysis in order to minimize the time of analysis, in this work 0 to 80 was used to account for the whole fermentation time.

The second common section for all pipelines is the Data pre-processing section, in which first the funciont `FCA_cData` helps cleaning the data of the instrumental noise. The latter is performed by eliminating negative, zero, infinite and NaN valued events in the relevant channels. Function allows to keep track of the proportion of this events and to select and if one wishes take out of the analysis data sets with noise above a certain threshold. Measurements with more than 2% of noise were not taken into account for further analysis. In this work all fermentation experiments the range of experimental points used for analysis ranged from 187 to 376 online flow cytometry readings,with a mean number of events above 20000 events. The majority of the samples with high instrumental noise were detected on the batch phase, which was not relevant on the continuous culture analysis done with the online flow cytometry. The higher instrumental noise in the batch phase is probably due to intermittent suboptimal online dilutions due to the constant growth of the strains. However a high coverage was achieved in the experimental setups replicates. The experimental set coverage can be observed in the Figure 1.

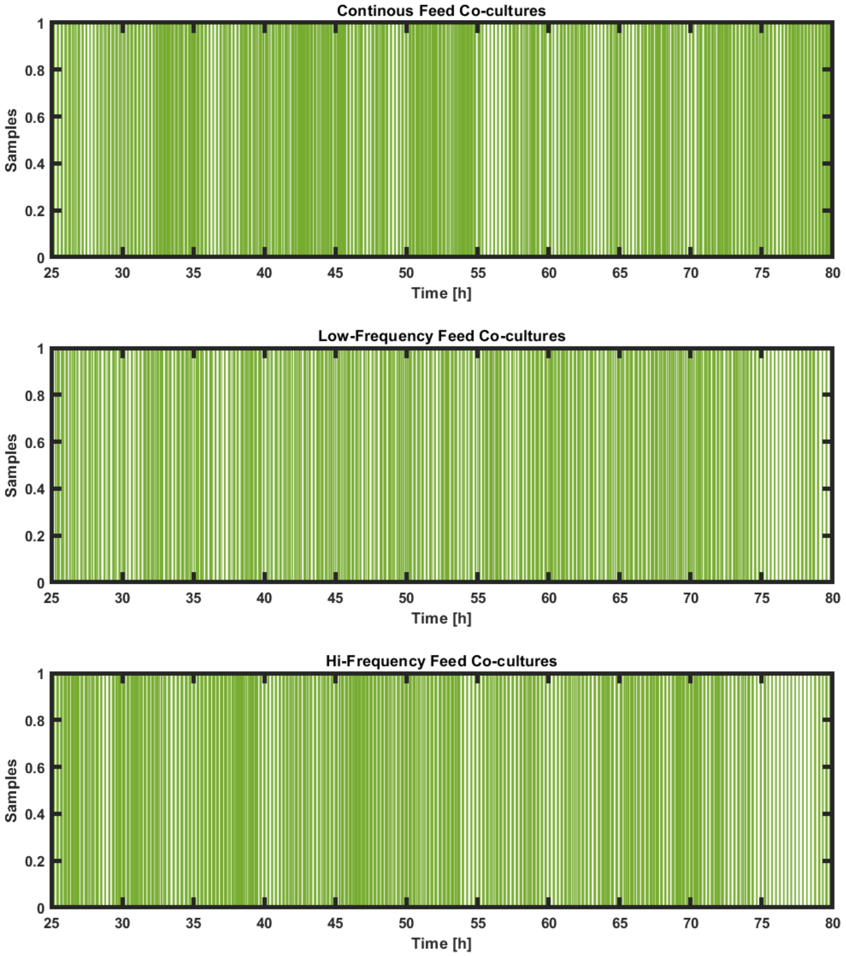


**Fig 1.** Measurement data set coverage for the Continuous, Low-frequency and High-frequency feed co-culture experiments from time 24 to 80.

After noise cleaning we quantified and eliminated Doublet events from each experimental measurement. The later was performed with the functions `FCA_FindDoublets` and the `FCA_Selection` functions. Briefly, on each measurement data set a linear function is aproximated for the the FSCH and FSCA channels, and the residuals for each event within this set calculated. The events with a person standarized residual value of 2 or more is deemed as a doublet. In this work none of the measurements had a fraction of doublets higher than 3% (Fig. 2).

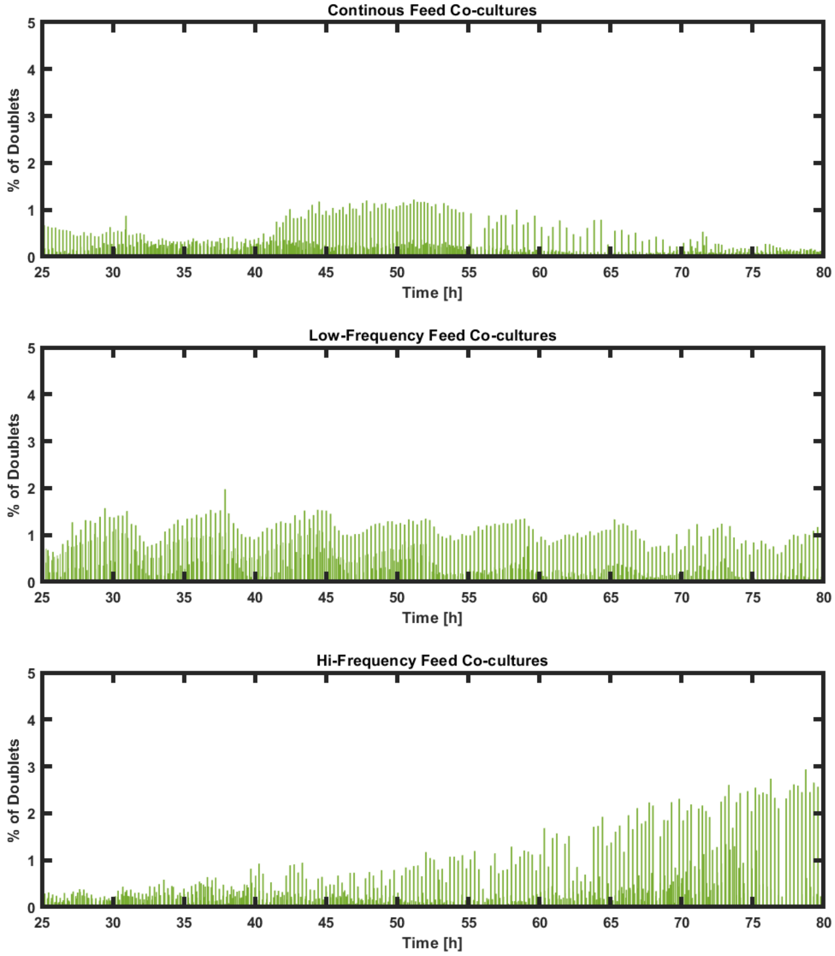


**Fig 2.** % of doublets found for each data set for the Continuous, Low-frequency and High-frequency feed co-culture replicate experiments from time 24 to 80.

After pre-treatment protocols, data is then processed for obtaining the statistical data. The function `FCA_Statistics`, is used to calculate the data range, mean, median, standard deviation, variance8 coefficient, variance, fano factor, interquartiles, and a probability density function. The probability density function used in this work was a normal distribution or a non-parametric kernel distribution.

After this initial analysis, the pipeline uses the probability density function and histogram of the FSCA channel to address to identify the events related to *S. cerevisiae* and *E. coli* according to a threshold set to be on the value of 5.3 log10(FSCA). The latter value was set based on the data obtained from the continuous cultivation of axenic cultures of the previously mentioned organisms (Fig. 3).

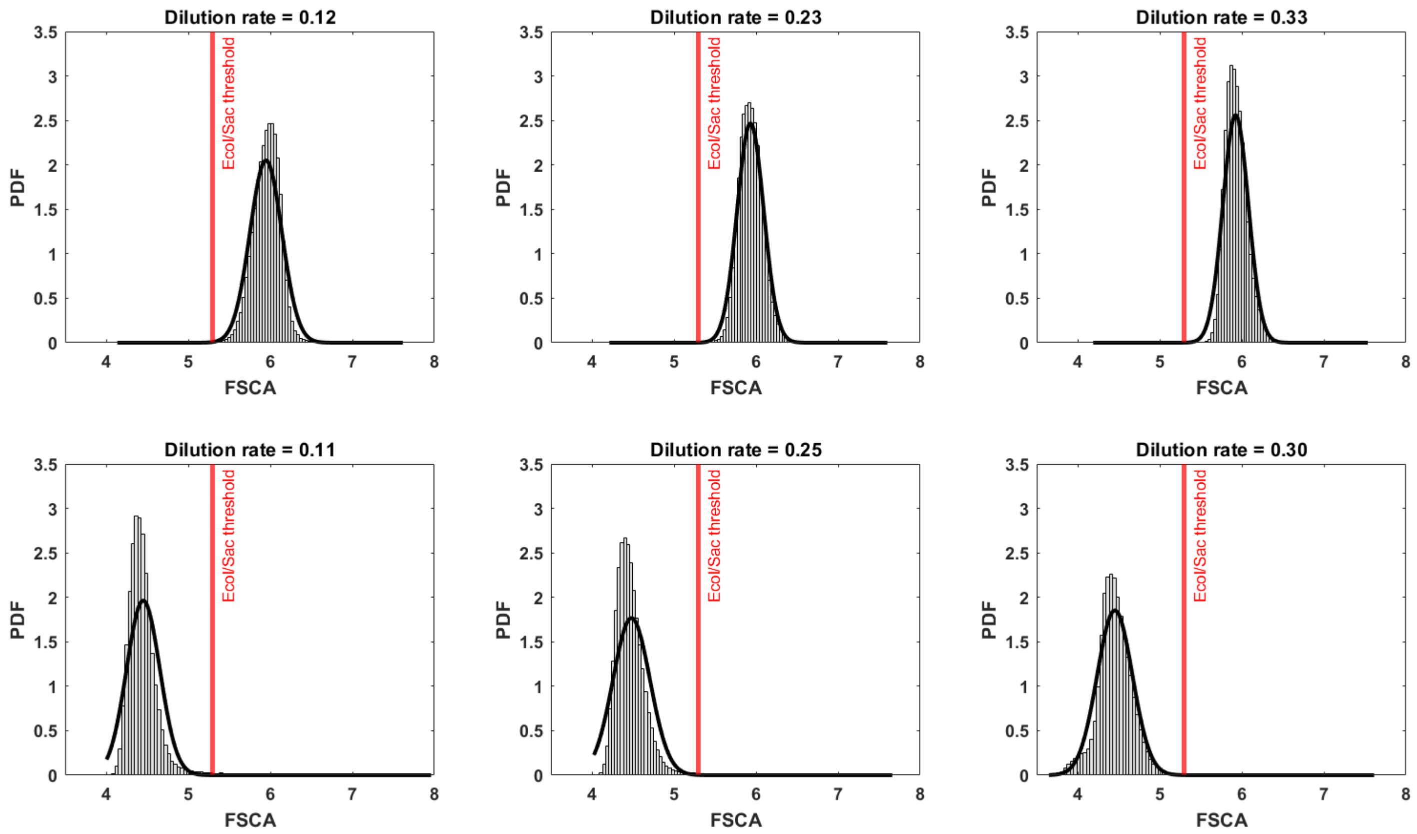


**Fig 3.** FSCA analysis for *S.cerevisiae* (up) and *E. coli* (down) and the selected threshold line on log10(FSCA) = 5.3 to address the clustering of each individual population during co-culture experiments.

The separation of the two populations at the above-metioned threshold is performed with the function `FCA_LineGate`, complemented with the use of the Selection function to write structures containing only the events of each organisms. The latter allow for quantification of their fraction, available at any experimental point. Finally, statistical analyses are made to each populations to address their individual mean, std, fano factor, within other statistical parameters.

In this particular pipiline example, after standar analyses, we decided to use a function called `FCA_GMValueClusters` that calculates the Grand Media and standard deviation for the all the experimental sets (4). With these values we quantify for each sample how many events are above the media plus three standard deviations. The results of these analyses be seen on figure 5.

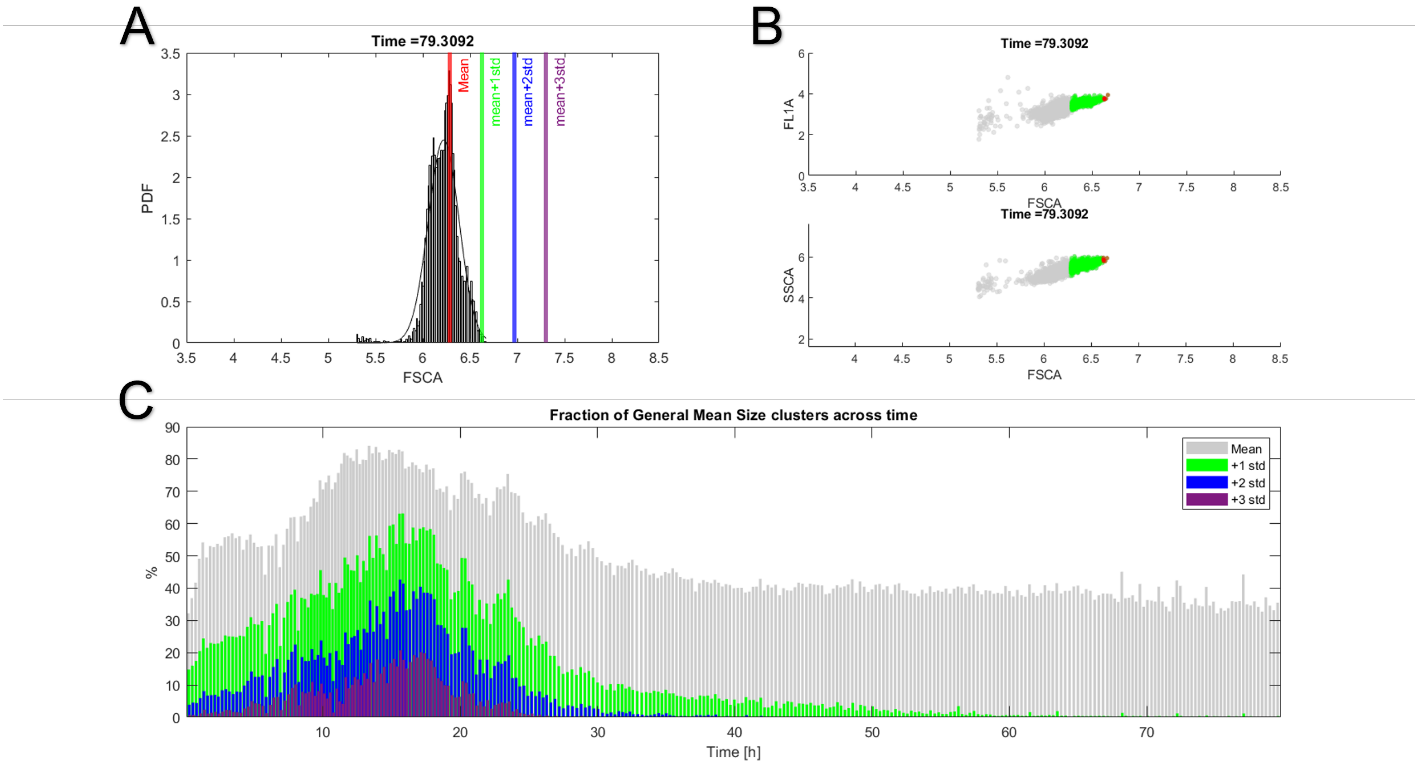


**Fig 4.** Example figure for the aggregate quantification protocol. A) Probability density function approximation and histogram compared to Grand mean and their standard deviations, for a sample at 79.3 h of the *S. cerevisiae* culsters of a high-frequency experiment. B) experimental scatter plots of data with colored fractions according to their FSCA distribution against grand media. C) Time evolution of each categorized population, the selected aggregates index of grand mean and 3 standard deviations is in purple

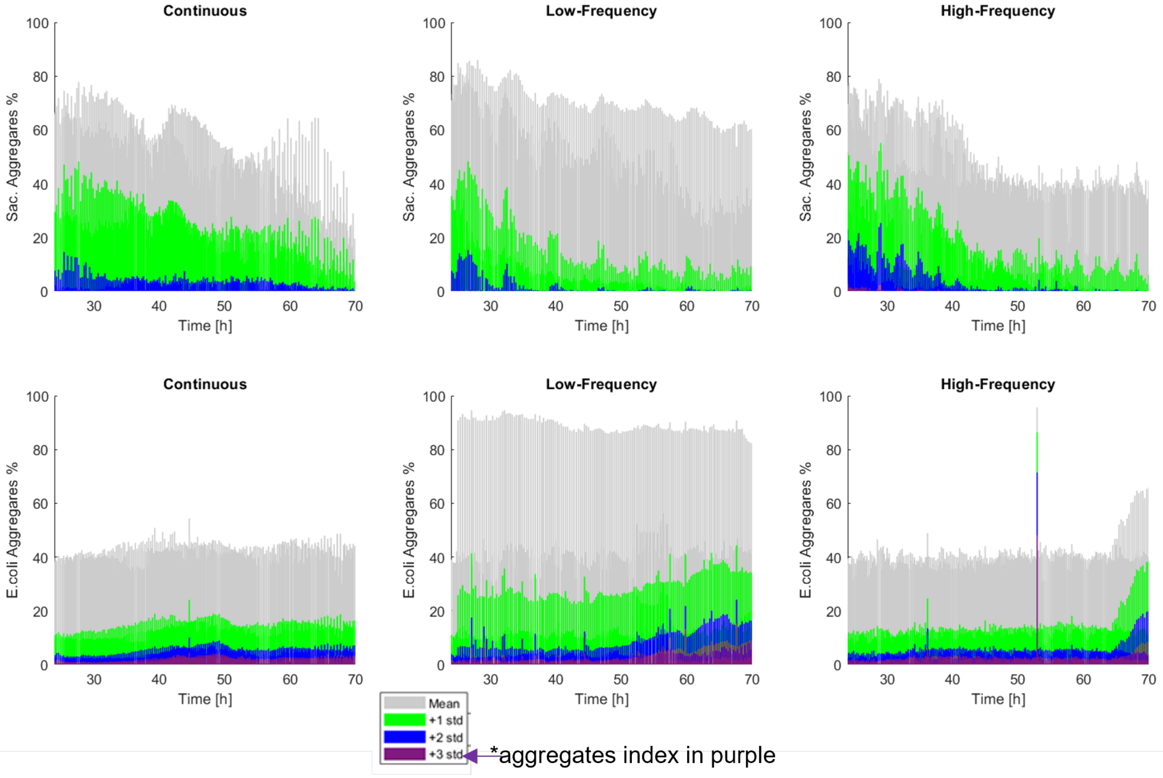


**Fig 5.** Time evolution of each categorized population against the grand media, aggregates selected by threshold in purple. Top row corresponds to the analysis for the three feeding profiles for the *S. cerevisiae* clusters. Bottom row corresponds to the analysis for the three feeding profiles for the *E. coli* clusters.

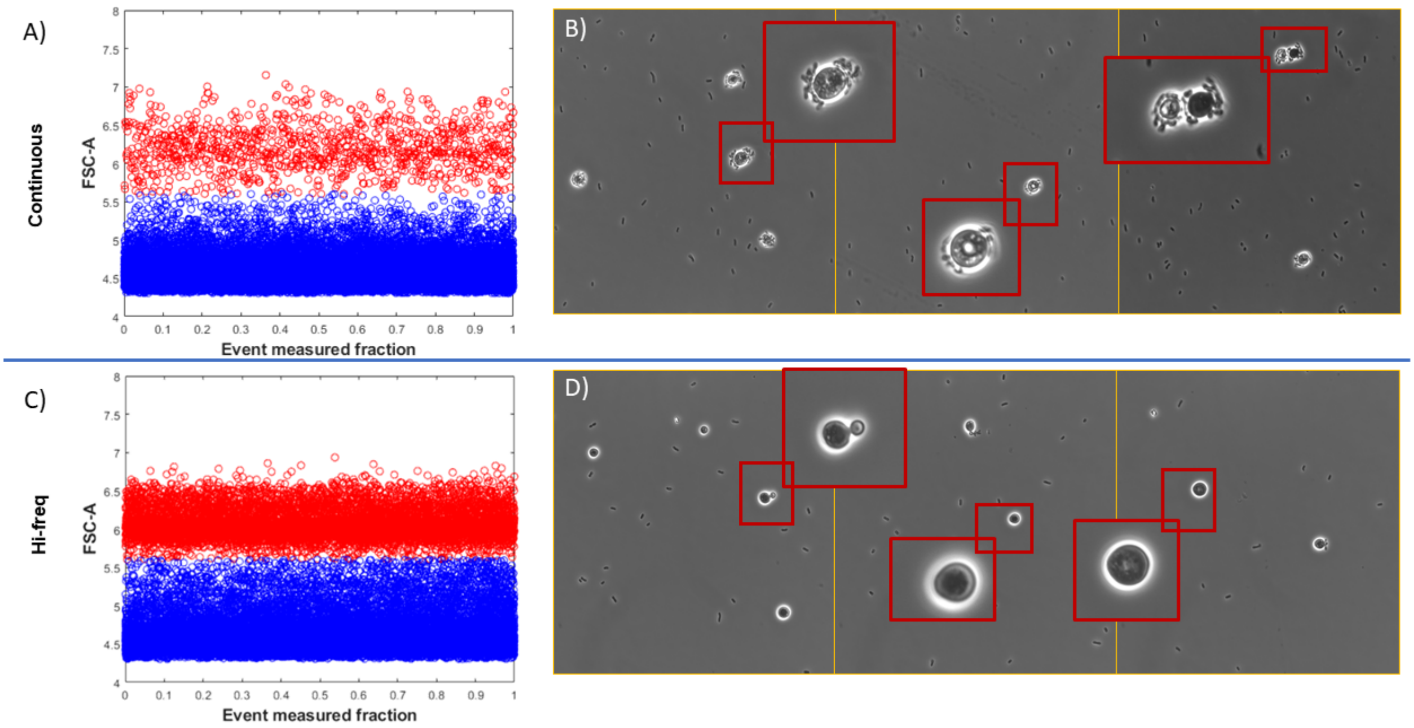


**Fig 6.** A) and B) Continuous cultivation FSC-A vs event measured fraction plot and micrography samples, respectively, at time 80 h. *E.coli* (blue) and *S.cerevisiae* (red). C) and D) Hi-frequency pulsed profile cultivation FSC-A vs SSC-A FC plot and micrography samples, respectively, at time 80 h. On all micrographies, example marked cells have been magnified.
