## Supplementary material for "Controlling microbial co-culture based on substrate pulsing can lead to stability through differential fitness advantages": S4_file

### Supplementary File 4: Continuous and Discontinuous culture supplementary figures

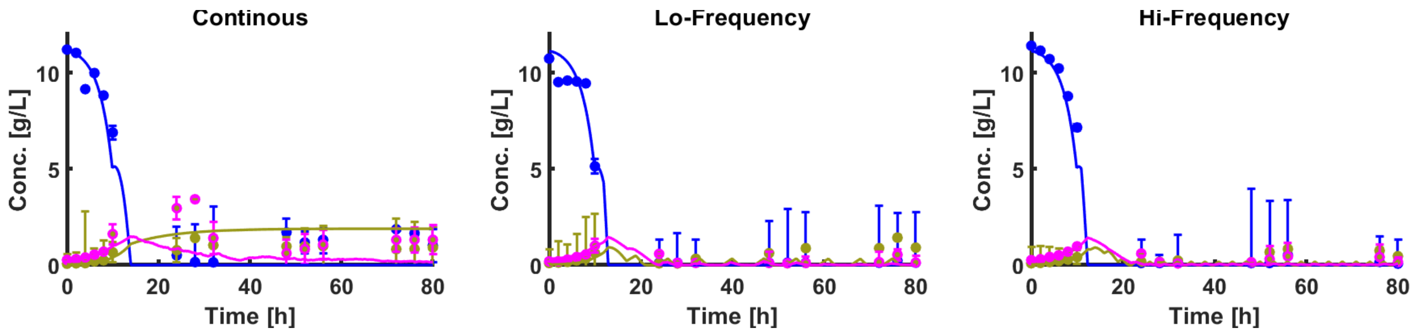


**Fig 1.** Measurement data set coverage for the Continuous, Low-frequency and High-frequency feed co-culture experiments from time 24 to 80.

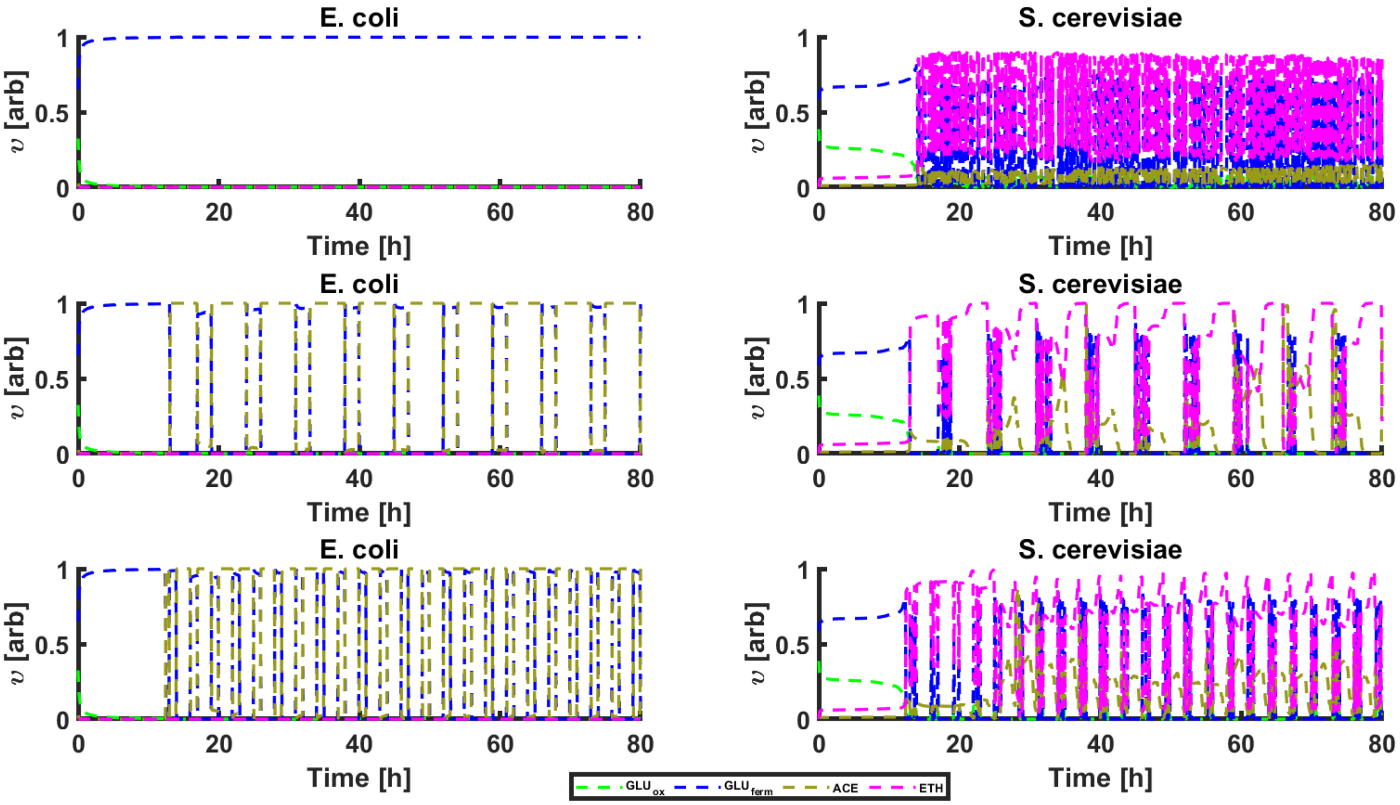


**Fig 2.** Cybernetic variable $v$ for *E. coli* and *S. cerevisiae* during the simulations made for Continuous culture (up), low frequency (middle) and high frequency (down) pulsing experiments.

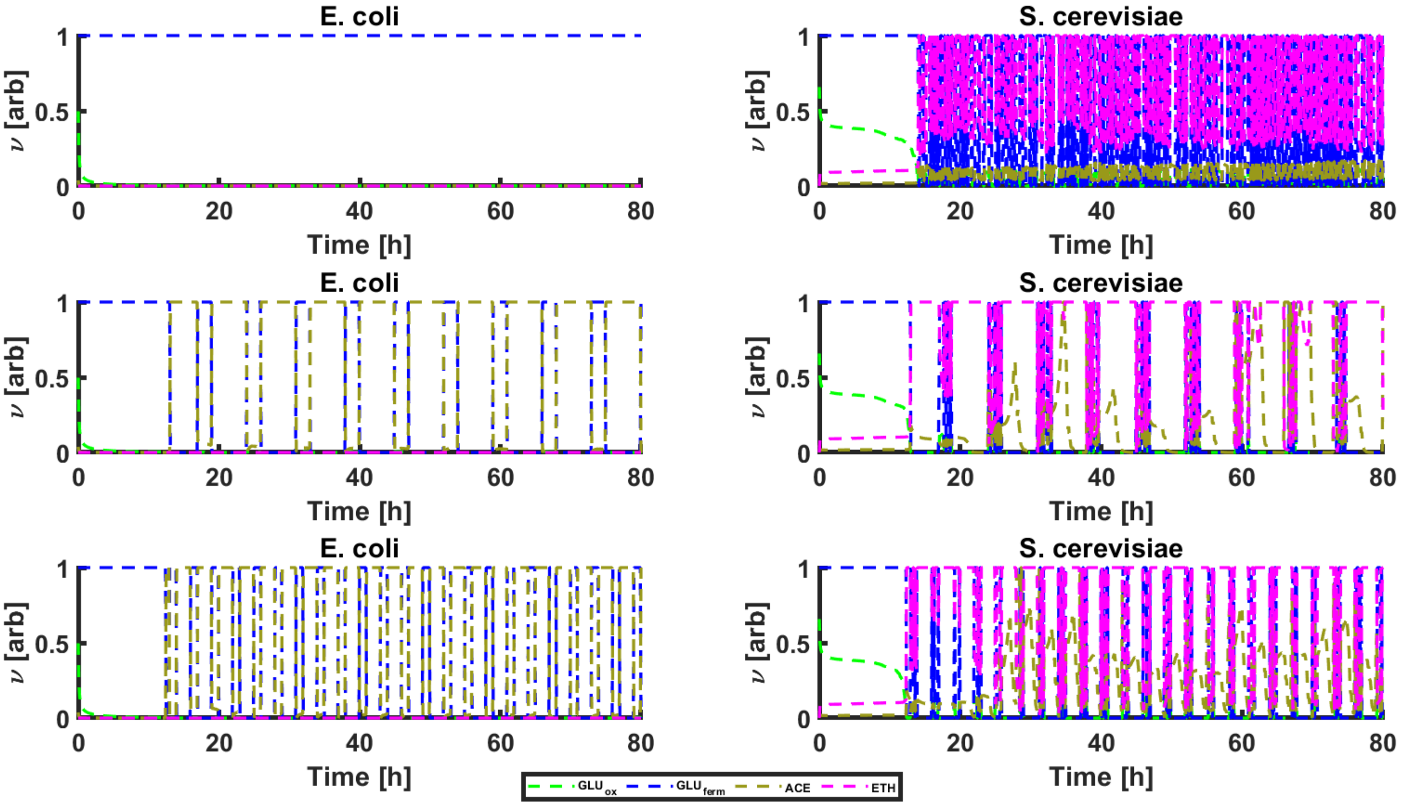


**Fig 3.** Cybernetic variable $\nu$ for *E. coli* and *S. cerevisiae* during the simulations made for Continuous culture (up), low frequency (middle) and high frequency (down) pulsing experiments.

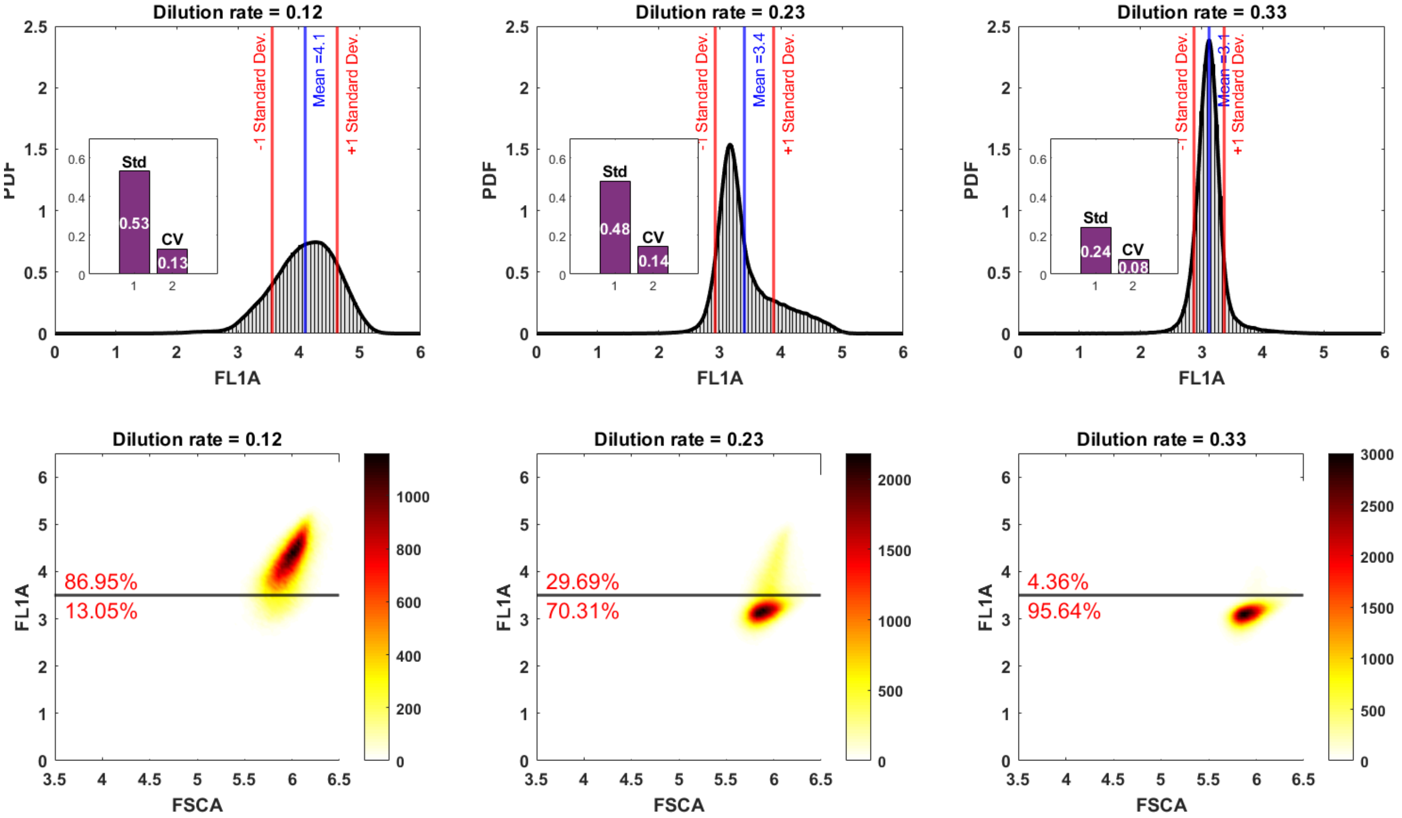


**Fig 4.** top row: FSCA Probability density function, mean, standard deviations and coefficient of variation for the axenic continuous cultures of *S. cerevisiae* at dilution rates of 0.12, 0.23 and 0.33 $h^{-1}$, respectively. bottom row: Flow citometry data for the triplicates at the different dilution rates and their $GFP^{+}$ and $GFP^{-}$ percentual calculations along the threshold at 3.5.

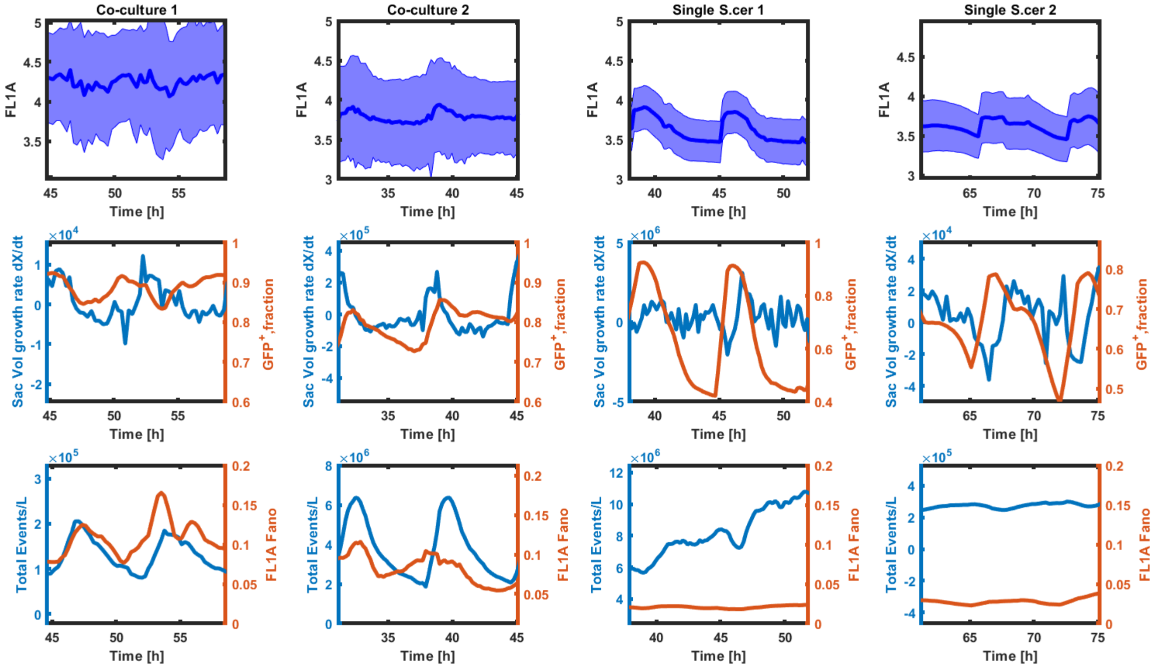


**Fig 5.** Time evolution of FL1, approximated growth rate, $GFP^+$ fraction, total events over liter and Fano factor across example cycles for the low-frequency feed regime experiments. Plotted values correspond for *S. cerevisiae* population in coculture and in single culture (duplicates)
